# Neural and Behavioral Dynamics of Acute Fentanyl Administration and Implications for Hazard Assessment of Novel Synthetic Opioids in Larval Zebrafish

**DOI:** 10.1101/2025.04.07.647579

**Authors:** Courtney Hillman, James Kearn, Gabrielle Wasser-Bennett, Maciej Trznadel, Matthew J. Winter, Matthew O. Parker

## Abstract

*Rationale* Synthetic opioids pose a significant public health risk due to their rapid synthesis and potentially lethal potency. New compounds are emerging continuously, meaning current testing platforms struggle to keep pace. Objectives Consequently, there is a critical need for simple, rapid, translatable models to provide a scalable screening platform to identify and hazard-assess emerging synthetic opioids, and test potential intervention strategies. Methods Here, we exposed 4 days post-fertilization (dpf) larval zebrafish to a range of concentrations of the prototypical class representative, fentanyl, to investigate behavioral and neural responses. Results Fentanyl caused low concentration hyperactivity, and high concentration hypolocomotion (sedation) which was reversed by the opioid antagonist naloxone. We confirmed predictive validity by replicating the behavioral responses with other class representatives (diacetylmorphine [heroin] and remifentanil). We also confirmed, pharmacologically, that low concentration hyperlocomotion was mediated by dopamine D2 receptors, replicating effects observed in mammals. Further mechanistic investigation using whole-brain *in vivo* imaging revealed disrupted connectivity in opioid-related circuits, such as the habenulae and dorsal thalamus, alongside novel pathways, including circuits associated with the pineal gland, torus semicircularis and eminentia granularis, potentially highlighting previously uncharacterized sensory and cerebellar neuronal networks. Conclusions These findings support the use of the larval zebrafish as a scalable model for assessment of synthetic opioids to provide novel insights into opioid-induced behaviors and mechanisms of action that may aid strategies in the growing challenge of interventive treatments for synthetic opioid intoxication.

**Impact Statement:** Larval zebrafish reveal neurobehavioral pathways affected by synthetic opioids, offering a valuable tool for rapid hazard assessment.

## Introduction

Opioids are a critical part of the arsenal to combat pain. However, opioid misuse in the United States of America (USA) has reached epidemic levels, with over 153 million opioid prescriptions and nearly 50,000 related deaths in 2019 (National Institute on Drug Abuse, 2021; Substance Abuse and Mental Health Services Administration, 2021). Other countries are starting to show a similar trend. In the United Kingdom (UK), for example, 5.6 million opioid prescriptions were dispensed between 2017 and 2018 with reports that approximately 15% of prescription users become long-term opioid abusers (Faria et al., 2018).

There is a growing prevalence of illicit heroin and other ‘street’ opioids containing novel synthetic opioids as adulterants (Giné et al., 2014 (United States Drug Enforcement Administration, 2024). For example, 73% of illicit fentanyl, a synthetic opioid with 50-100 times the potency of heroin, is consumed unknowingly (Rodda et al., 2017; World Health Organization, 2021). Fentanyl is implicated in over half of annual opioid overdose fatalities (Arendt, 2021) and requires higher doses of naloxone to reverse due to its high lipophilicity and enhanced blood-brain barrier (BBB) penetration (Eastlack et al., 2020; Nóbrega & Dinis-Oliveira, 2018; Shafi et al., 2020; Sutcliffe et al., 2022). Synthetic derivatives like ohmefentanyl and carfentanil pose even greater overdose risks due to their extreme potency (Chen et al., 2001; Swanson et al., 2017; H. Wang et al., 1991).

Addressing the evolving opioid epidemic requires models to assess the potency and risks of synthetic opioids. Zebrafish (*Danio rerio*) are a powerful and cost-effective alternative to traditional mammalian models, offering high fecundity, suitability for rapid, large-scale *in vivo* analyses and unique opportunities for whole-brain, cell-level imaging *in vivo* (Hillman et al., 2024a; A. Stewart et al., 2011; A. M. Stewart et al., 2012, 2014; A. M. Stewart & Kalueff, 2014; Vanwalleghem et al., 2018). Using larvae ≤ 4 days post-fertilization [dpf]) also benefits from the reduced ethical constraints due to being unprotected under the Animals (Scientific Procedures) Act (1986). Despite recent studies showing behavioral responses to fentanyl (Kirla et al., 2021; Pesavento et al., 2022; B. Wang et al., 2022) and autonomic dysregulation with a novel respiratory depression assay (Zaig et al., 2021), no research has evaluated opioid-induced behavioral effects in 4 dpf larvae or explored their impact on brain function using whole brain functional imaging.

This study evaluates the prototypical synthetic opioid fentanyl in 4 dpf larvae using a novel behavioral assay suitable for the rapid assessment of opioid effects. To investigate the mechanistic basis of the observed behavior, we then analyzed whole-brain functional responses to fentanyl using *in vivo* neural imaging. Fentanyl produced consistent concentration-dependent behavioral responses. Sedative and hyperactive fentanyl-induced changes in neural activity and whole-brain functional connectivity *in vivo* provided insights into the neurobiological pathways affected by exposure. Together, these approaches allow us to assess both the behavioral and neural impacts of synthetic opioids in larval zebrafish, offering the basis for a scalable and ethically preferable *in vivo* model for rapid opioid hazard prediction.

## Results

### Concentration-dependent behavioral effects of fentanyl

A robust, high-throughput, cost-effective model is needed to rapidly assess the effects of novel synthetic opioids, and larval zebrafish (≤ 4 dpf) provide an ethically advantageous solution. Previous research using older (5 dpf) larval zebrafish in the light/dark assay has revealed fentanyl concentration-dependent (0.001 μM – 13 μM) hypolocomotion (but not full sedation) in the dark phase (Colwill & Creton, 2011; Kirla et al., 2021; Pesavento et al., 2022; B. Wang et al., 2022). Therefore, we first carried out a full comprehensive analysis of the light/dark behavioral effects of fentanyl in 4 dpf zebrafish larvae to a wide range of concentrations of fentanyl (0.05 μM – 50 μM), using our validated light/dark locomotor assay (Hillman et al., 2024a, 2024b) (**Fig. 1**). This revealed a biphasic locomotor response characterized by hyperactivity at lower concentrations (0.05 µM), progressing to hypoactivity (5, 20 and 50 µM) and complete immobility with increasing concentrations (**Fig. 1A**).

**Fig. 1.**
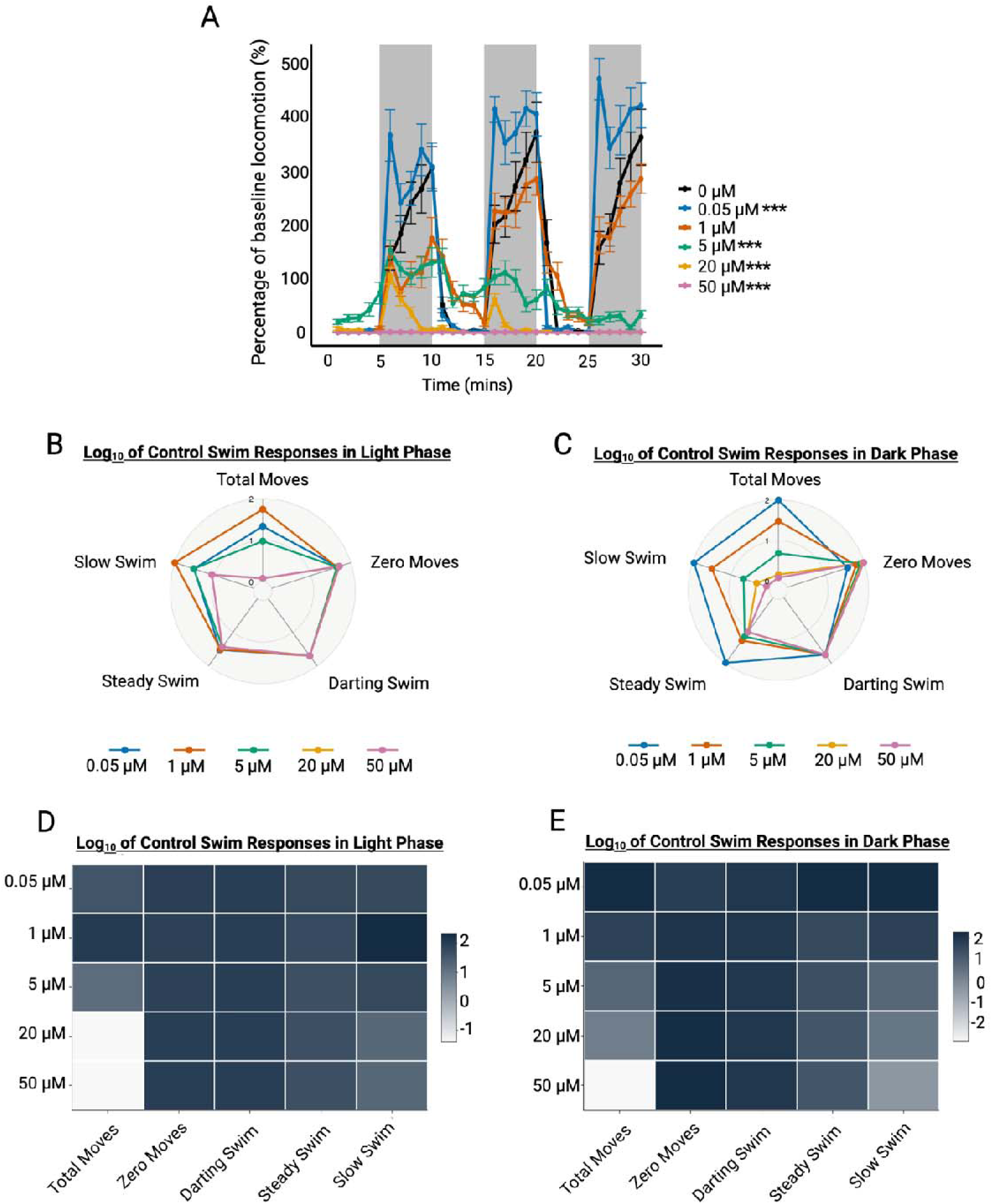
Concentration-dependent effects of fentanyl on 4 dpf larval behavior in the light/dark assay. (**A**) Analysis represented as mean percentage of baseline movement over the 30-min recording period following exposure to a range of fentanyl concentrations (0.05 μM – 50 μM). Larvae display low concentration hyperactivity followed by concentration-dependent hypolocomotion in the dark phase (shaded in grey) compared to vehicle-exposed controls confirmed with a generalized linear mixed model (GLMM) with a gamma distribution on dark phase locomotion (β = 5.43, SE = 0.12, z = 46.50). The likelihood ratio test (LRT) against the null model (χ^2^(5) = 86.93, *p* < 0.001) confirmed including concentration-dependent effects on behavior significantly improved the fit of the model. This was further characterized by a significant increase in dark phase locomotion following exposure to a low concentration (0.05 μM; Dunn’s multiple comparison test: Z = -5.09, *p* < 0.001) followed by a significant decrease with 5 μM (Dunn’s multiple comparison test: Z = 9.01, *p* < 0.001. A time-dependent effect was assessed with a GLMM with gamma distribution (β = 0.0046, SE = 0.0028, z = 47.41) and was not statistically significant when tested with an LRT against the null model (χ^2^(1) = 2.81, *p* = 0.09). (**B** and **D**) Light phase swim phenotype responses shown as a radar chart and heat map. The larvae exposed to the higher concentrations of fentanyl (5 μM, 20 μM and 50 μM) had decreased ‘total moves’ compared to vehicle-controls (*p* = 0.0275, *p* < 0.0001, *p* < 0.0001). The two highest concentrations (20 μM and 50 μM) also had significantly decreased ‘zero moves’ compared to vehicle-controls (*p* < 0.0001, for both). (**C** and **E**) show the dark phase swim phenotype responses as a radar chart and heat map. 0.05 μM fentanyl exposed larvae resulted in increased ‘total moves’ compared to vehicle-controls (*p* < 0.0001) followed by concentration-dependent reduction in ‘total moves.’ ‘Zero moves’, ‘steady swim’ and ‘slow swim’ were also affected by fentanyl exposure with significantly decreased ‘steady swim’ and ‘slow swim’ compared to vehicle-controls with the highest tested concentrations (5 μM, 20 μM and 50 μM). Significantly increased ‘zero moves’ was seen with all three of the highest concentrations (5 μM (*p* = 0.02), 20 μM (*p* = 0.014) and 50 μM (*p* = 0.0002)). Data in panel **A** is presented as mean ± standard error of the mean (SEM). In panels **B**, **C**, **D** and **E**, data were presented as mean. (*n* = 16, observed power = 0.997). *Note: *** = Significantly different dark phase locomotion compared to vehicle-exposed control at the p<0.001 level*.

To further explore these concentration-dependent behavioral responses, we analyzed discrete swim phenotypes (**Fig. 1B-E**). Overall, dark phase ‘total moves,’ ‘zero moves’, ‘steady swim’ and ‘slow swim’ responses were significantly affected by fentanyl exposure with significantly increased ‘total moves’, ‘steady swim’ and ‘slow swim’ responses with hyperactive (0.05 µM) fentanyl. Comparatively, these phenotypes were significantly decreased with 5 µM, 20 µM and 50 µM. These concentrations also resulted in significantly increased ‘zero moves’ compared to vehicle-controls.

### Behavioral Effects of Fentanyl are Likely Mediated by μ-Opioid Receptors

Following these light/dark locomotor experiments, we confirmed bioavailability of fentanyl using LC/MS-MS (see supplementary materials) within our exposure window. Next, to determine whether the observed locomotor responses were a result of opioid receptor activation, we co-administered 20 μM naloxone, a non-specific opioid receptor antagonist, with 5 μM fentanyl. Reductions in dark phase locomotion by 5 μM fentanyl were fully rescued with co-administration of naloxone, with no significant difference seen between the co-administration and vehicle-control groups (**Fig. 2**). Additionally, naloxone alone had no significant impact on dark phase locomotor responses. Given naloxone’s pronounced affinity for the μ-opioid receptor (MOR), these results strongly suggest that fentanyl-induced behavioral changes in zebrafish larvae are mediated predominantly through MOR activation.

**Fig. 2.**
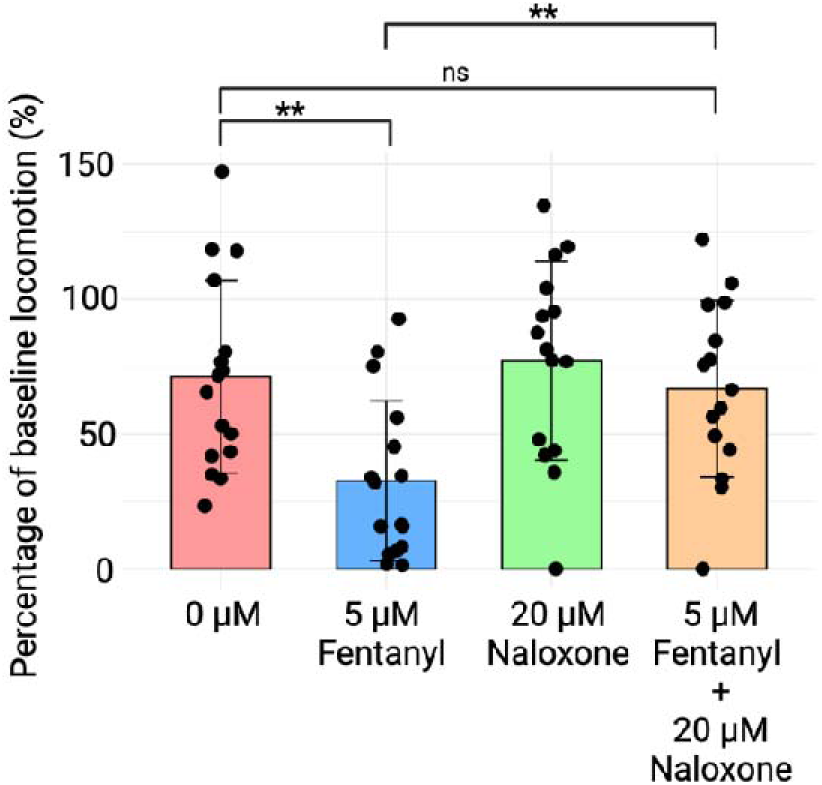
Opioid receptor pharmacology regulating locomotor response in larval zebrafish. The averaged dark phase locomotor response in 4 dpf larval zebrafish exposed to 5 μM fentanyl, 20 μM naloxone and co-administration of both. These data show fentanyl significantly reducing dark phase locomotion compared to vehicle-controls (*p* = 0.011), which was significantly reversed with co-administration of 20 μM naloxone (*p* = 0.984). Naloxone alone had no significant effect (*p* = 0.961). Normalized data are presented as mean ± SEM. Circles indicate individual data points for each zebrafish. (*n* = 16, observed power = 0.9496). *Note: ** = Significantly different dark phase locomotion compared to control at the p<0.01 level*.

### Low-Concentration Fentanyl-Induced Hyperlocomotion is Mediated by Dopamine

Low concentrations of fentanyl elicit hypermobility in rodents via dopamine receptor-mediated rewarding effects (David et al., 2002; Le Merrer et al., 2009; Le Merrer et al., 2007). We therefore hypothesized the hyperactivity observed here may result from dopaminergic signaling. Sulpiride, a selective dopamine D2 receptor antagonist (Nakazato et al., 1998), diminishes the rewarding effects of opioids in rodents as evidenced by its reversing of opioid-induced hypermobility (Morgenstern & Fink, 1985). The hyperlocomotor response to 0.05 μM fentanyl was fully rescued by 100 μM sulpiride (**Fig. 3**). Importantly, sulpiride alone had no significant effect on behavioral outcomes. These results provide evidence of dopaminergic involvement in fentanyl-induced responses and the first demonstration of the neuropharmacological translational potential of the larval zebrafish model for opioid rewarding effects.

**Fig. 3.**
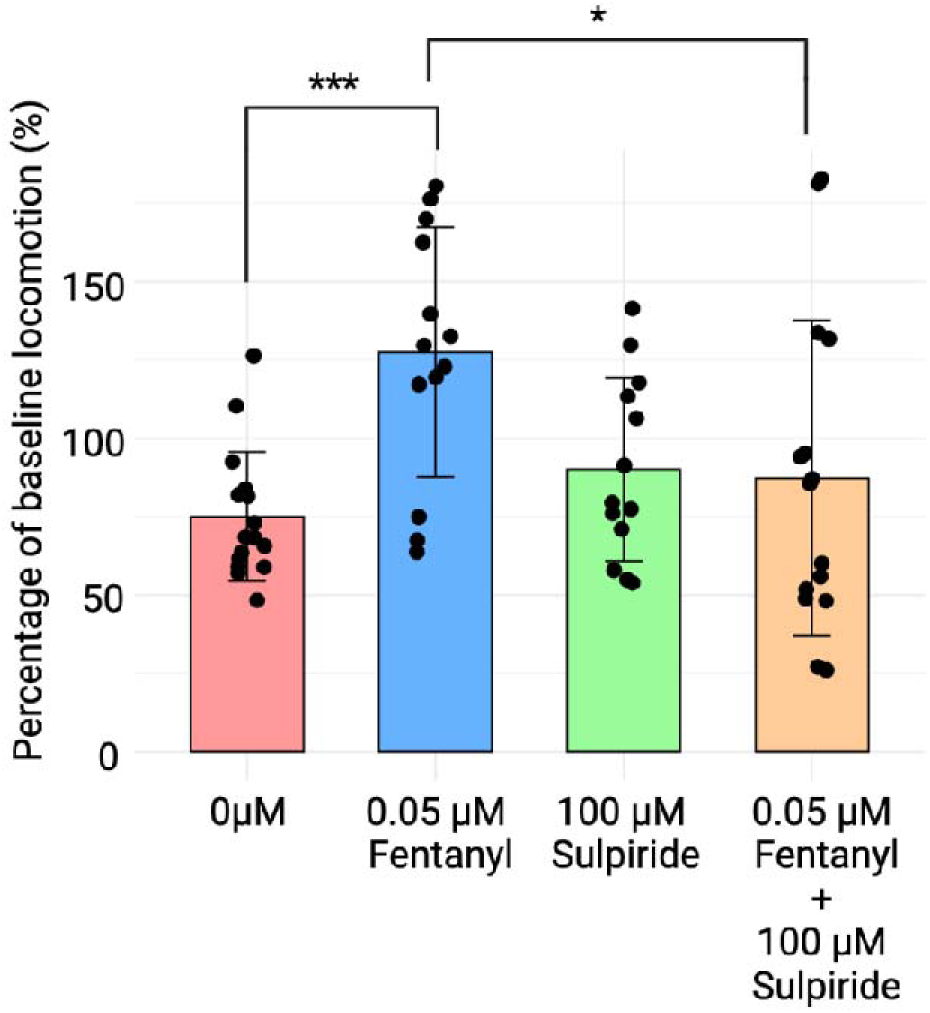
Dopamine D2 receptor antagonism reverses fentanyl-induced larval hyperactivity. The averaged dark phase locomotor response in 4 dpf larval zebrafish exposed to 0.05 μM fentanyl, 100 μM sulpiride and co-administration of both showing full rescue of hyperlocomotion with sulpiride (one-way ANOVA: F_(3, 53)_ = 5.26, *p* = 0.00299). Fentanyl significantly increasing dark phase locomotion compared to vehicle-controls (*p* = 0.001). This was reversed with co-administration of 100 μM sulpiride (*p* = 0.027). Normalized data are presented as mean ± SEM. Circles indicate individual data points for each zebrafish. (*n* = 16, observed power = 0.9574). *Note: * indicates significantly different dark phase locomotion*.

### General predictive validity of locomotor assay for opioid detection

Next, we aimed to assess the generalized predictive validity for detecting opioids of varying potencies by exposing 4 dpf larvae to other opioids (diacetylmorphine [heroin] and remifentanil) (Choi et al., 2008; Hill et al., 2020; Komatsu et al., 2007; Koyyalagunta, 2006; Thompson & Rowbotham, 1996) (**Fig. 4**). Low concentrations of heroin produced comparable locomotor responses to that of fentanyl, for example, inducing dark phase hyperactivity followed by concentration-dependent hypolocomotion. Specifically, heroin caused a significant increase in dark phase locomotion at 0.05 μM compared to vehicle controls (**Fig. 4A**), while remifentanil did not show this effect at the same concentration (**Fig. 4F**). Remifentanil demonstrated a time-dependent effect, with its initial marked inhibitory effect reducing in magnitude by the final dark phase. In contrast, fentanyl caused a time-dependent decrease in dark phase locomotion. At concentrations above 50 μM, heroin exposure resulted in increased movement relative to control during the light phase (**Fig. 4A**). Granular analysis using radar charts and heat maps (**Fig. 4B-D**) illustrates the significant increase in light phase locomotion, in particular ‘total moves’, a response not observed with the other opioids.

**Fig. 4.**
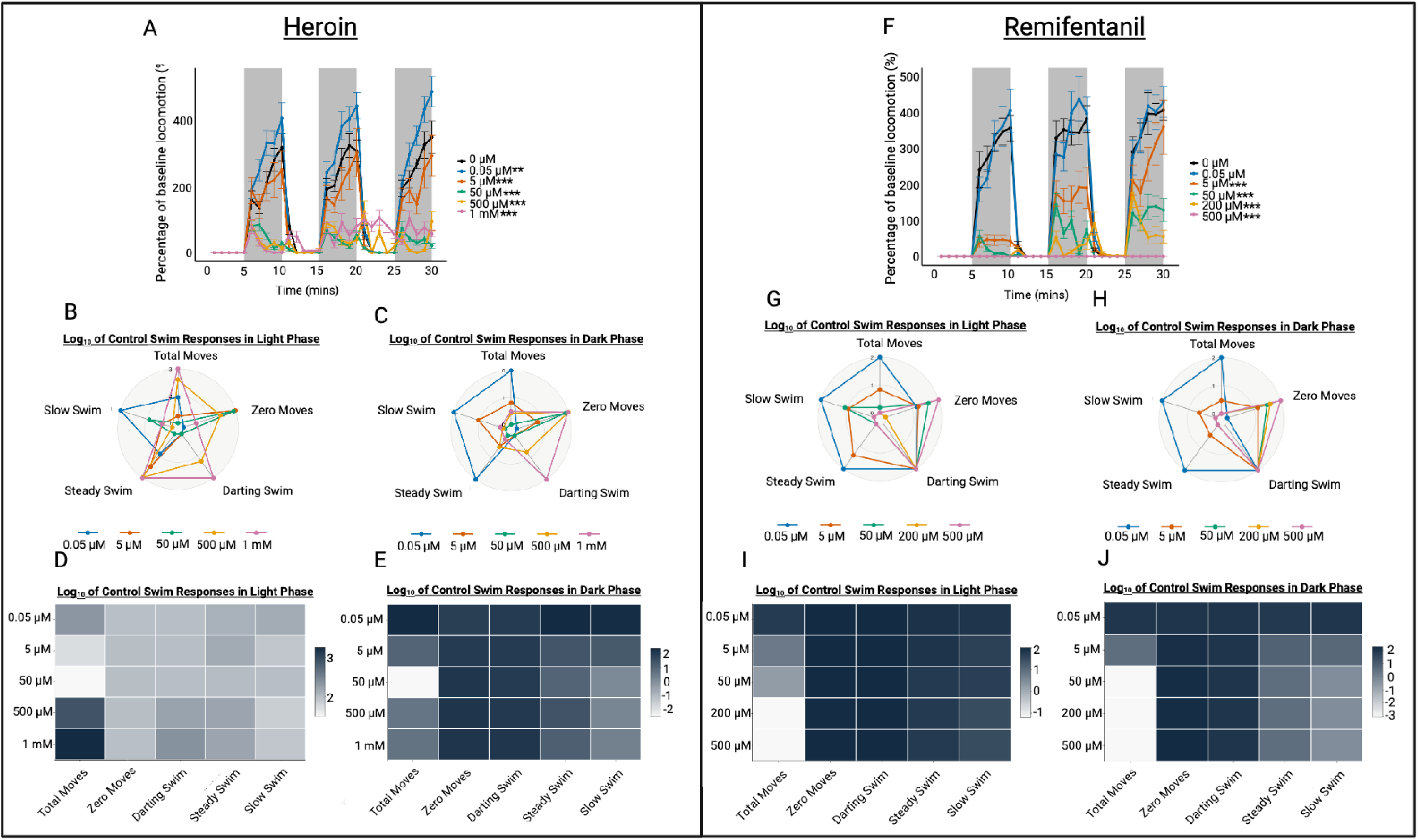
Concentration-dependent effects of heroin and remifentanil on 4 dpf larval behavior in the light/dark assay. (**A**) The time series analysis is represented as mean percentage of baseline movement for each minute over the 30-min recording period following exposure to a range of heroin concentrations (0.05 mM – 1 mM). Larvae showed dark phase hyperactivity with low concentrations of heroin, followed by concentration-dependent hypolocomotion compared to vehicle-controls (GLMM with gamma distribution: β = 5.46, SE = 0.12, z = 43.93; LRT vs. null model: χ^2^(5) = 89.6, *p* < 0.001). The three highest tested concentrations (50 μM – 1 mM) resulted in a loss of distinctive light/dark locomotion which was not seen with fentanyl. (**B** and **C**) Light phase swim phenotype responses shown as a radar chart and heat map. These data show that the top tested concentrations (500 μM and 1 mM) showed a loss of light/dark response across the time series analysis (*p* = 0.034 and *p* = 0.022, respectively). These concentrations also showed increased ‘total moves’ compared to vehicle-controls (*p* < 0.0001, respectively). (**D** and **E**) Further analysis of the dark phase data showed decreased ‘total moves’ responses with 500 μM and 1 mM (*p* < 0.0001, respectively). There were also significant increases in ‘total moves’, ‘slow swim’ and ‘steady swim’ moves observed at the lowest tested concentration (0.05 μM) (*p* = 0.0001, *p* = 0.0007 and *p* = 0.0003, respectively). (*n* = 16, observed power = 0.925). **(F**) The same time series data are shown as in panel (**A**) but following exposure to a range of remifentanil concentrations (0.05 μM – 500 μM). These larvae showed dark phase concentration-dependent hypolocomotion compared to the vehicle-controls with a time-dependent effect (GLMM with gamma distribution (time effect): β = 0.03, SE = 0.003, z = 10.04; LRT χ^2^(1) = 94.37, *p* < 0.001). (**G** and **H**) Light phase swim phenotypic responses shown as a radar chart and heat map. Here, the ‘total moves’ (*p* < 0.0001), ‘zero moves’ (*p* < 0.0001), steady swim’ (*p* = 0.0001) and ‘slow swim’ (*p* < 0.0001) responses were all significantly affected by remifentanil exposure. (**I** and **J**) Further analysis of the dark phase phenotype data showed all swim responses to be significantly affected with remifentanil exposure (*p* < 0.0001, respectively), except for darting movements. (*n* = 16, observed power = 0.983). Data in panels **A** and **F** are presented as mean ± standard error of the mean (SEM). In panels **B** – **E** and **G – J**, data were presented as mean. *Note: ** = Significantly different dark phase locomotion compared to control at the p<0.01 level*.

To further explore the predictive validity of the assay, we produced concentration-response curves comparing the respective potencies of the tested opioids (**Fig. 5**). All three compounds produced predictable sigmoidal curves with dark phase locomotion decreasing as concentration increases (relative potencies [IC_50_]: fentanyl > remifentanil > heroin). Remifentanil has a reduced potency relative to fentanyl (Choi et al., 2008; Komatsu et al., 2007; Thompson & Rowbotham, 1996), possibly due to remifentanil’s metabolic pathway (Duthie, 1998). Although it has a similar mode of action to the other opioids, remifentanil is unique because of its esterase metabolism, causing an ultrashort duration of action (Duthie, 1998). As such, when used as an analgesic or anesthetic in a clinical setting there is a need for a continuous infusion, which may explain why remifentanil’s behavioral response appeared to be short-lived, suggesting a similar metabolic profile and response as seen in higher organisms.

**Fig. 5.**
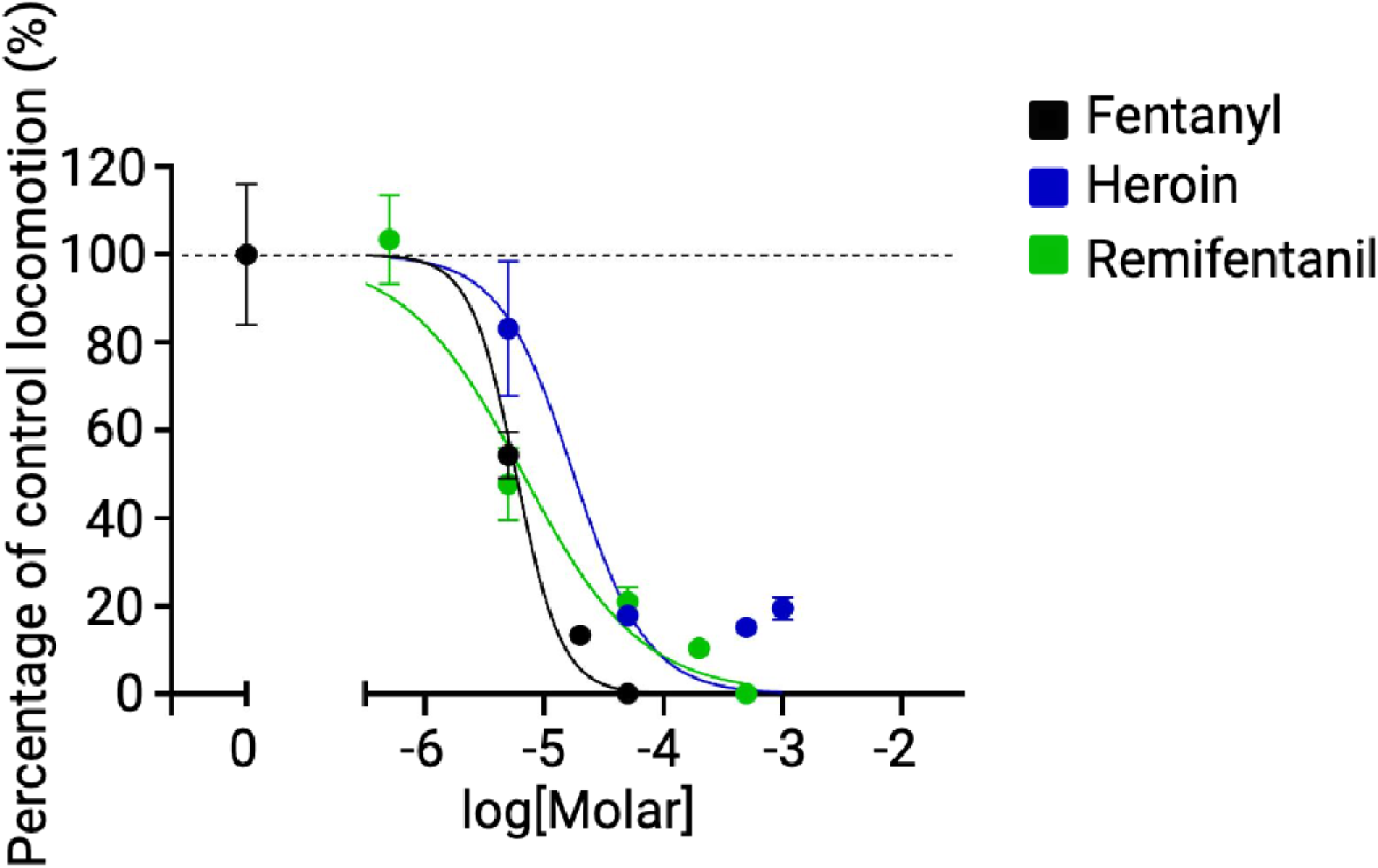
Opioid concentration-response curves. The opioid concentration-response curves for fentanyl, heroin and remifentanil excluding the hyperactive concentrations displayed as percentage of vehicle-control dark phase responses and logarithmic transformation of the molar concentration using a non-linear regression. The curves suggest a relative potency as follows: fentanyl > remifentanil > heroin, with calculated LogIC_50_ values of -5.256 (5.54 μM) for fentanyl, -5.176 (6.67 μM) for remifentanil and - 4.761 (17.3 μM) for heroin. Data are represented as mean ± SEM with *n* = 16 for each concentration. The data point at log[Molar] = 0 represents the vehicle-control response for reference.

### Whole-brain imaging reveals novel neural circuits affected by synthetic opioids

The optical transparency of larval zebrafish affords us the unique capacity to visualize how fentanyl affects neural activity across various brain regions during administration. Combining our behavioral evaluation with neuronal investigations allows for validation of behavioral outcomes, combined with intricate analysis of potential neurological mechanisms. Using a functional neuroimaging approach (adapted from Winter et al. (2017, 2021), originally described by Vladimirov et al. (2014)), we observed qualitative differences in fluorescence intensity in specific brain regions between vehicle-treated controls and larvae exposed to hyperactive (0.05 μM) and sedative concentrations of fentanyl (50 μM) (**Fig. 6**). Sedative fentanyl significantly affected activity in regions associated with sensory and motor coordination, including the anterior commissure and eminentia granularis (Bae et al., 2009; Li et al., 2024; Pose-Méndez et al., 2023); areas involved in homeostatic and regulatory functions, such as the vagal ganglia, rostral hypothalamus and preoptic area (Higashijima et al., 2000; Machluf et al., 2011; Palieri et al., 2024; Tsuneoka & Funato, 2021); and the posterior tuberculum which is implicated in reward processing and motor control (Haehnel-Taguchi et al., 2018). All areas saw an increase in activity compared to vehicle-controls, except for the vagal ganglia and eminentia granularis, which were significantly reduced (**Fig. 6B**).

**Fig. 6.**
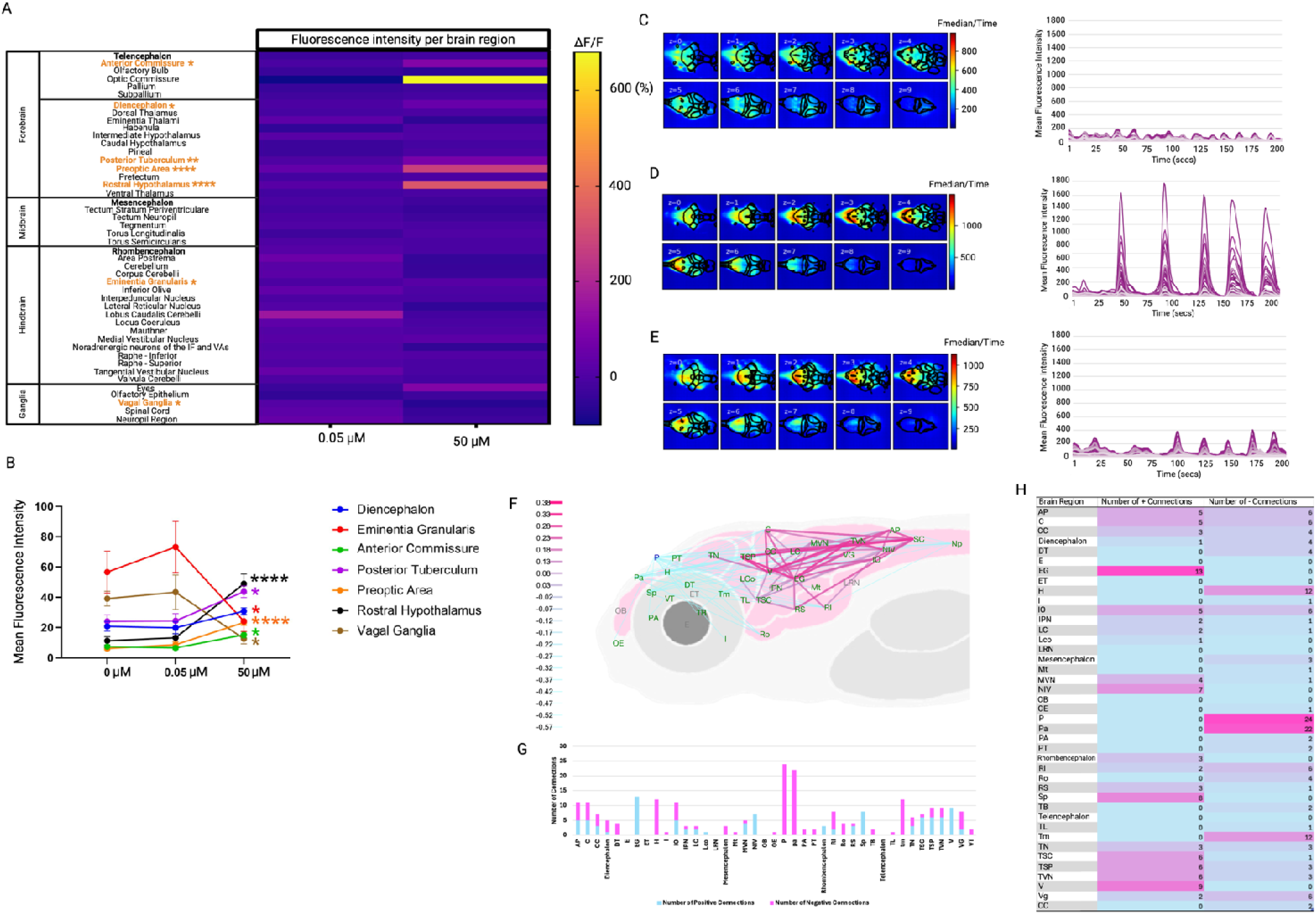
Functional 4-dimensional brain profiling and average functional connectivity across the brain in fentanyl-treated 4 dpf zebrafish larvae. (**A**) Summary of percentage change of fluorescence intensity data per brain region of interest (ROI) compared to vehicle-controls. These data are presented as the average baseline-corrected fluorescence intensity (Δf/f = (f_1_ – f_0_)/f_o_ *100, where f_1_ = peak fluorescence intensity and f_0_ = vehicle-control fluorescence intensity within each ROI expressed as percentage increase over the same ROI from the corresponding vehicle control group. The results show a general decrease in activity at a high concentration (sedative) of fentanyl compared to vehicle-controls and increase in activity at low concentrations (hyperactive) (*n* = 7 larvae per group). *Note: * and orange text = Significantly different dark phase locomotion compared to control at the p<0.01 level.* (**B**) Concentration-response mean fluorescence for the ROIs with post-hoc significance identified (one-way ANOVA: F_(2, 18)_ = 3.693, *p* = 0.05). (**C**) The median fluorescence intensity projections across time, for each z-plane (0-9 ventral to dorsal surface) for vehicle-controls, (**D**) for 0.05 μM fentanyl and (**E**) for 50 μM fentanyl. To the left of each image, a color scale representing the fluorescence intensity range averaged over the experimental duration is shown for reference. The results clearly show the differences in fluorescence intensity within specific brain regions in each z-slice. (**F**) Functional connectome showing pairwise connections that had *r*-values that were significantly different between vehicle-controls and 0.05 μM fentanyl exposed larvae. Nodes connected with a pink line represent significant increases and blue as decreases in the *r* value compared with controls. (**G** and **H**) display the total number of significant positive or negative connections identified for each brain region in stacked bar chart format (**G**) and tabulated (**F**). The highest number of increased *r* values were seen associated with the eminentia granularis, valvulae cerebelli and noradrenergic neurons of IF and VAs with the highest number of decreased *r* values associated with the pineal gland and pallium. Data in (**B**) are shown as connections which produced a different mean *r* value compared with the equivalent connection as a mean of all vehicle-control larvae tested. Abbreviated brain regions are as follows: area postrema (AP); cerebellum (C); corpus cerebelli (CC); dorsal thalamus (DT); eyes (E); eminentia granularis (EG); eminentia thalami (ET); habenulae (H); intermediate hypothalamus (I); inferior olive (IO); interpeduncular nucleus (IPN); lobus caudalis cerebelli (LC); locus coeruleus (Lco); lateral reticular nucleus (LRN); mauthner (Mt); medial vestibular nucleus (MVN); noradrenergic neurons of the interfascicular and vagal areas (NIV); spinal cord neurophil (Np); olfactory bulb (OB); olfactory epithelium (OE); pineal (P); preoptic area (PA); pallium (pa); pretectum (PT); raphe inferior (RI); rostral hypothalamus (Ro); raphe superior (RS); spinal cord (SC); subpallium (Sp); posterior tuberculum (TB); torus longitudinaris (TL); tegmentum (tm); tectum neurophil (TN); torus semicircularis (TSC); tectum stratum periventriculare (TSP); tangential vestibular nucleus (TVN); valvula cerebelli (V); vagal ganglia (VG); ventral thalamus (VT). *Note: Positions of regions approximate to allow spacing of nodes and optimal visualization of connectivity*.

To further explore the effects on neural activity, we conducted functional connectivity analyses, comparing the temporal profiles of activity between pairs of regions of interest (ROIs) in fentanyl-treated versus vehicle-control larvae. In the context of neural imaging, functional connectivity refers to the relationship between temporal activity profiles of anatomically separate regions as an indicator of the level of communication between these regions (Vladimirov et al., 2014). Such relationships can be measured as correlations which can be compared between drug-treated and untreated brain states. Hyperactive (0.05 μM) fentanyl exposure revealed significant differences in connections involving the eminentia granularis, noradrenergic neurons of IF and Vas, valvulae cerebelli, pineal gland, habenulae, tegmentum and pallium (**Fig. 6**). A substantial proportion of the significant connections identified were negative, including a strong reduction in functional connectivity between the habenula and pretectum as well as between the habenula and the dorsal thalamus between fentanyl treated and control larvae. The habenulae are involved in negative reward processing and aversive behaviors (Mondoloni et al., 2022), whereas the pretectum and dorsal thalamus are involved in visual and sensory processing (Rees & Roberts, 1993; Usrey & Alitto, 2015). This suggests altered connectivity between regions involved in aversion and withdrawal as well as altered pain processing. We also observed a significant decrease in functional connectivity between the pineal gland and cerebellum, suggesting an impact on autonomic processing and motor function.

Several of the pathways identified were consistent with characterized circuits involved in brain opioid responsiveness. However, the analysis also revealed several previously uncharacterized pathways which may be involved in the rewarding circuitry of opioids, including a unique increase in functional connectivity between the torus semicircularis and eminentia granularis. These are pathways involved in sensory processing and cerebellar functioning and are not commonly identified in opioid-related circuitry (Folgueira et al., 2020; Privat et al., 2019). We also observed previously uncharacterized opioid-related pathways involving the pineal gland, including reduced connectivity with the tangential vestibular nucleus and the cerebellum between treated and untreated animals.

## Discussion

With the continual emergence of novel synthetic opioids as illicit drugs, there is an urgent need for a rapid, high-throughput, and translationally relevant models to predict opioid potency and health risks as soon as new compounds are identified. In this study, we defined the 4 dpf larval zebrafish light/dark locomotor assay responsiveness to fentanyl and confirmed the opioid-specific effects by co-administering the antagonist naloxone. We then reported successful reversal of low concentration hyperactivity by co-administering the dopamine D2 receptor antagonist, sulpiride. Next, we validated the model’s translational and comparative power by assessing responses across two further opioids, remifentanil and heroin. Finally, we exploited the optical transparency of larvae to map the brain’s functional response to fentanyl exposure *in vivo* for the first time using light-sheet microscopy, identifying specific brain regions affected by fentanyl exposure. These findings collectively support the utility of the 4 dpf zebrafish model for rapid assessment of synthetic opioid effects on behavior and brain activity. They also highlight potential novel regions of interest (ROIs) for opioid action in the brain, providing insights that could inform understanding of the mechanisms of action and possible behavioral effects of newly emerging opioids of abuse.

To date, no published studies have mapped the effects of synthetic opioids on whole-brain function *in vivo* while correlating these effects with behavioral responses in larval zebrafish. Using transgenic zebrafish expressing genetically-encoded Ca^2^L reporters and light sheet microscopy, we identified several brain regions significantly affected by fentanyl, including areas involved in motor control (e.g., anterior commissure, eminentia granularis) (Bae et al., 2009; Li et al., 2024; Pose-Méndez et al., 2023), regulatory functions (e.g., vagal ganglia, rostral hypothalamus, preotic area) (Higashijima et al., 2000; Machluf et al., 2011; Palieri et al., 2024; Tsuneoka & Funato, 2021), and reward processing (e.g., posterior tuberculum) (Haehnel-Taguchi et al., 2018). A concentration-dependent response was observed, with sedative doses causing notable decreases in fluorescence in the eminentia granularis and vagal ganglia, reflecting opioid-induced disruption of sensory-motor coordination and autonomic functions (Bae et al., 2009; Pose-Méndez et al., 2023). Functional connectivity analysis revealed disrupted pathways, particularly involving the habenulae, a key region in sensory processing and emotional regulation (Ables et al., 2023; Fore et al., 2018; Proulx et al., 2014; Rees & Roberts, 1993), with reduced connectivity to the dorsal thalamus and pretectum, suggesting impaired aversive processing and reward sensitivity (Lawson et al., 2014; Metzger et al., 2021; Stephenson-Jones et al., 2012). Significant alterations in dopaminergic pathways, with diminished connectivity in the tegmentum and pineal gland, indicated possible disruptions in circadian, sensory-motor, and reward circuits (Gorwood, 2008; Upadhyay et al., 2010). Novel connectivity changes, including those between the torus semicircularis and eminentia granularis, suggested dysregulated sensory-motor integration underlying hyperactive behaviors (Edwards & Kelley, 2001; Knogler et al., 2017).

Fentanyl disrupts a broad network of neural circuits, from opioid-sensitive areas to downstream regions critical for respiratory and motor functions, offering insights into potential mechanisms behind opioid-induced dysregulation. Whole-brain evaluation of fentanyl exposure successfully identified its availability in the larval zebrafish brain as well as highlighting novel ROIs, providing avenues for future research to identify novel opioid mechanisms of action.

We acknowledge several limitations of our study. First, we estimate the uptake and bioavailability of fentanyl using whole-body homogenates (**Supplementary Materials**), which does not specifically confirm neuronal availability. Although the light sheet microscopy data provides additional clarity, there remains a possibility that the reduced locomotion in the larvae may be the result of the presence of fentanyl in the exposure solution rather than the pharmacological effects of the drug at its target tissue. For example, fentanyl or other substances could have noxious or aversive effects that affect locomotion, although the light sheet data and bioavailability measurements suggest this is unlikely (Wisenden et al., 2022). There is also a possibility that the exposure solutions may degrade over time, which may account for some of the time-dependent effects that were seen. This would however require further investigation to confirm. Additionally, there is limited data available on adult zebrafish for comparison, which could impact the interpretation of age-specific effects. The study may offer an incomplete hazard assessment model for novel synthetic opioids, as it does not capture all dimensions of opioid toxicity, such as long-term effects or specific toxicological endpoints like respiratory depression. However, some of these limitations may be mitigated through a multi-model approach, potentially incorporating older larvae and adults which can be used to study other opioid-induced phenomenon including respiratory depression (Zaig et al., 2021). Finally, while functional connectivity analysis provides valuable insights into network dynamics, the correlational nature of this approach limits our ability to infer direct causality. Experimental approaches, such as optogenetics or chemogenetics, may help to validate and expand upon these findings in future studies.

This study offers new insights into the neuronal and behavioral effects of synthetic opioids, reporting the first data on behavioral impacts in 4 dpf larvae, supporting their use as tool for rapid screening of emerging opioid drugs of abuse in a 3Rs-prioritized, translationally relevant, vertebrate model. Such insights could inform the development of targeted treatments for synthetic opioid-induced disruptions, and a scalable and ethical tool for estimation of relative potency of emerging synthetic opioids of health concern.

## Materials and methods

### Animal Husbandry

Wild-type (WT) and transgenic (Tg) zebrafish (*Danio rerio*) used in this study originated from in-house breeding. For behavioral studies, WT AB-strain adult zebrafish were housed in mixed-sex groups (approximately 50:50 male to female ratio) with 8-10 fish per 2.8L tank or up to 20 per 6L tank, maintained in a recirculating water system (Aquaneering, Fairfield Conrolec Ltd., Grimsby, UK). Water temperature was regulated between 25-27°C, with a pH of 8.4 (±0.4), under a 14-hour light, 10-hour dark cycle, where lights turned on at 8:30 AM. From 5 dpf onwards, larvae were fed ZM-100 fry food (Zebrafish Management Ltd., Winchester, UK) until reaching adulthood, at which point their diet was transitioned to flake food and live brine shrimp, fed three times daily (once per day on weekends). The larvae used in these experiments were derived from over 50 independent breeding sessions, achieved through pair breeding and the addition of substrate to the home tanks. Further details on breeding protocols are provided in Hillman *et al*. (2024b). Larvae were housed in a translucent incubator at 28.5°C until they were transferred to a small countertop incubator at 28°C for behavioural testing. Upon completion of testing, the larvae were euthanized using Aqua-Sed (Aqua-SedTM, Vetark, Winchester, UK) according to the manufacturer’s instructions, followed by maceration. All experiments were approved by the Animal Welfare and Ethical Review Boards of the University of Surrey.

Light sheet microscopy and bioanalysis were performed using Tg zebrafish expressing the pan-neuronal calcium sensor *elavl3 (elavl3:GCaMP6)* and in-house WT strain broodstock, respectively, both of which were housed in flow-through aquaria (Techniplast, VA, Italy) in The University of Exeter Aquatic Resource Centre (ARC). Adult zebrafish broodstock were reared under optimal conditions for spawning (28+/- 1°C; 12-hour light: 12-hour dark cycle, with 20-minute dusk–dawn transition periods). Animals were cultured in mains tap water, which was filtered by reverse osmosis (RO; Osmonics E625 with cellulose membranes; GE Water and Process Technologies, Trevose, PA, USA), and then reconstituted with commercial marine salts (Tropic Marin), calcium chloride and sodium bicarbonate to create standardized synthetic freshwater (final salt concentrations provided a conductivity of 1000 μS). This culture water was also aerated and heated to 28 ± 2 °C in a reservoir before it was supplied to the stock holding tanks within a recirculating aquaculture system. Adult fish were fed twice daily with a mixed diet of commercial pellet food and cultured artemia. Culture water was routinely monitored for pH, temperature, conductivity, alkalinity, ammonia, nitrate and nitrite, which were maintained within acceptable limits of U.S. EPA guidelines (U.S. EPA, 1986). The same water and conditions were used to culture embryo-larvae and for the dilution of solutions for drug exposures (referred to as exposure water) prior to functional brain imaging. Spawning was triggered at the onset of light by introducing a spawning substrate. Fertilized eggs were collected, rinsed thoroughly, and then transferred to Petri dishes containing culture water. Embryos were kept under conditions consistent with the adult fish cultures until reaching 4 dpf. All procedures were reviewed and approved by the Animal Welfare and Ethical Review Body (AWERB) at the University of Exeter and carried out under UK Home Office regulations.

### Behavioral Apparatus

All behavioral tests were performed between 09:00 – 18:00 (Mon-Fri) using Zantiks MWP units (Zantiks Ltd., Cambridge, UK). Video recording was undertaken from an overhead position, allowing for real-time monitoring of fish behavior.

### Light/Dark Behavioral Assay

A detailed protocol for the light/dark behavioral assay is available in Hillman et al. (2024b), and its implementation is demonstrated in Hillman et al. (2024a). Briefly, larvae were transferred into a bench-top incubator in the behavioral testing room at 8:30 AM, set to 28°C for a 30-minute acclimation period. For testing, 225 µL of Petri dish water (with same composition and pH as culture water described above) and a randomly selected larva was transferred to each well of a 48-well plate using modified pipette tips (approximately 2 mm trimmed from the end of a 1000 µL pipette tip). Larvae were chosen at random from different Petri dishes. The well plate was then placed in the Zantiks MWP unit, where the overhead lighting was switched on, and larvae were allowed to acclimate for 30 minutes. Following this acclimation period, the light/dark tracking protocol was initiated for baseline measurements, consisting of three light-to-dark transitions, each phase lasting five minutes, with the light intensity set to 350 lx (white light). The temperature of the MWP unit was maintained at 28.02°C ± 0.07 throughout the experiment. After the baseline recording, the well plate was removed, and drug treatments (see *Opioids and Concentration Range Tested* for more information) were administered by adding 25 µL of the prepared solution from a 96-well plate to achieve the desired final concentration. The well plate was then returned to the MWP unit, and the larvae were tracked through further light/dark cycles post-treatment.

### Opioids and Concentration Range Tested

Table 1 provides an overview of all opioid drugs (fentanyl, heroin and remifentanil) used in the present study. All three opioids were tested at five concentrations plus vehicle DMSO 0.5% (V/V) (Hoyberghs et al., 2021)). The opioid receptor antagonist naloxone was tested at one concentration only. Fentanyl has previously been tested using the larval light/dark assay (Kirla et al., 2021; Pesavento et al., 2022; Sales Cadena et al., 2021; B. Wang et al., 2022; Zaig et al., 2021), however, no analysis with 4 dpf larvae using the tested opioids has been published to date. Therefore, the references were used as a concentration prediction only with all concentrations ranges determined through preliminary dose-response findings in our lab. This also applied for the naloxone and sulpiride concentrations tested.

**Table 1.** Compound overview including the supplier, catalogue number, concentration range tested, reported LogP value and references.

| Compound | Supplier | Catalogue Number | Concentrations [μM (mg/L)] | LogP | References |
| --- | --- | --- | --- | --- | --- |
| Diacetylmorphine (Heroin) | Sigma-Aldrich | H159 | 0.05 – 1000 (0.018 – 369.41) | 1.58 | National Center for Biotechnology Information, 2024f |
| Fentanyl citrate salt (Fentanyl) | Sigma-Aldrich | F3886 | 0.05 – 50 (0.026 – 26.43) | 4.05 | Cooman et al., 2021; Kirla et al., 2021; National Center for Biotechnology Information, 2024e; Pesavento et al., 2022; B. Wang et al., 2022; Zaig et al., 2021 |
| Tricaine methanesulfonate (MS-222) | Sigma-Aldrich | E10521 | 7654.3 (2.0) | 2.12 | National Center for Biotechnology Information, 2024b |
| Naloxone | Sigma-Aldrich | 1453005 | 20 (6.55) | 2.09 | National Center for Biotechnology Information, 2024a; Zaig et al., 2021 |
| Remifentanyl hydrochloride (Remifentanyl) | Sigma-Aldrich | Y0001892 | 0.05 – 500 (0.02 – 206.46) | 1.45 | National Center for Biotechnology Information, 2024c |
| (±)-Sulpiride | Sigma-Aldrich | S8010 | 100 (34.14) | 0.6 | National Center for Biotechnology Information, 2024d |

All drugs were made up as stock solutions in 100% DMSO and stored frozen where applicable until further dilution with water (with the same composition as the exposure water) to obtain the final working solutions and a final DMSO concentration of 0.5% (V/V).

### Light/Dark Assay Data Analysis

Raw data files were imported into a custom R script (R Core Team, 2024) available on <u>OSF</u>, where they were normalized relative to baseline values. Normalization was achieved by calculating the percentage movement of each larva relative to the overall baseline mean response. Outliers were identified and excluded if they exceeded the absolute deviation from the median, following the method described by Leys et al. (2013). To assess data distribution, residuals from null linear mixed models were plotted as QQ-plots and histograms. Since the data did not follow a normal distribution, a zero-inflated gamma (ZIG) generalized linear mixed model (GLMM) was employed. The dependent variable was percentage movement, while independent variables included light phase, drug concentration, and time, analyzed using the glmmTMB package in R. Fish ID was included as a random effect. The ZIG-GLMM model was selected due to the prevalence of 0% movement data, and its suitability for analyzing skewed continuous variables (Zuur & Ieno, 2021). Model fit was evaluated using likelihood ratio tests (LRTs) by comparing the full model with a null model. The same approach was applied for the dark phase data separately. Additionally, a Kruskal-Wallis test followed by Dunn’s post-hoc test was used to analyze the effect of drug concentration on locomotion during the dark phase. Significance thresholds were set at *p*\*\*\* < 0.001, *p\*\** < 0.01, *p\** < 0.05.

Radar charts and heat maps were generated using the custom R processing script, analyzing movement data as a percentage of baseline as darting (>2000%/s), steady swimming (690– 2000%/s), slow swimming (0.1–690%/s) or absence of movement (0%/s). To account for zero responses, a value of one was added to each observation prior to percentage of baseline calculations. Movement endpoints included total movements, darting movements, steady swimming, slow swimming, and zero movements. Mean values for these endpoints were calculated for the control groups. Movement percentages for each drug concentration were compared to vehicle-control groups and logarithmically transformed (base 10). Light and dark phase swimming behavior distributions were assessed using QQ-plots and the Shapiro-Wilk test, revealing normal distributions for dark phase data and non-normal for light phase data. Consequently, a one-way analysis of variance (ANOVA) and Dunnett’s post-hoc test were used to analyze dark phase phenotypic movement data. Light phase data was assessed with Kruskal-Wallis tests and Dunn’s multiple comparison post-hoc tests.

### Concentration-response curves

The concentration-response curves were created using the raw data, following standard outlier removal as described in Light/Dark Assay Data Analysis. This raw data was then exported into Microsoft Excel (Microsoft Excel, 2024) where the offset function was used to calculate the per five-minute movement. The mean dark phase response was calculated and then each concentration of opioid was calculated as a percentage of control dark phase response as follows: ((fish X concentration X)/average vehicle-control dark phase response*100). For more information see the excel file in OSF. A logarithmic conversion was performed on the molar concentration using log_10_ and all non-hyperactive concentrations were plotted in GraphPad Prism 10 (GraphPad Software, version 10.2.3, 2024) using a non-linear regression (log(inhibitor) vs. normalized response – Variable slope).

### Bioanalysis of larval internal (whole body) compound concentration

Fentanyl uptake analysis was carried out at the University of Exeter following the basic protocol outlined in Winter *et al*. (2021). Briefly, groups of four WT larvae were exposed to four concentrations of fentanyl (2, 5, 10, and 20 µM) as well as vehicle-controls for 30 minutes. After exposure, larvae were euthanized in 2 mg/mL tricaine methansulfonate (MS-222) at pH 7.5 and transferred along with 300 µL of the exposure solution to a 96-well filter plate (Multiscreen HTS BV 1.2 µm, Millipore, UK). The residual solution was removed under vacuum, and the larvae were washed with twelve 300 µL volumes of 0.5 mg/mL MS-222 in rig water (pH 7.5), followed by vacuum removal of the wash solution. The larvae were then transferred to a 96-well plate with 300 µL of water, a 4 mm stainless steel ball bearing, and 300 µL of a 50 nM internal standard in acetonitrile. The samples were homogenized using a Genogrinder (Spex, Metuchen, NJ, USA) at 1000 strokes per minute for 3 minutes. After homogenization, 900 µL of MS-grade water was added, resulting in a final mixture of 80:20 water to acetonitrile containing 10 nM of the internal standard (IS). The plate was agitated to mix the contents, then centrifuged at 4000 rpm for 30 minutes. A 900 µL aliquot of the supernatant was transferred to a separate 96-well plate for analysis by liquid chromatography-tandem mass spectrometry (LC-MS/MS). Similarly, 300 µL samples of the exposure solution were processed in the same manner for comparison.

Analysis of samples was performed using a Surveyor MS Pump Plus high-performance liquid chromatography (HPLC) system with an HTC PAL autosampler, coupled to a TSQ Vantage triple quadrupole mass spectrometer with a heated electrospray ionization (HESI II) source (ThermoFisher Scientific, Hemel Hempstead, UK). Chromatographic separation was achieved using a reversed-phase C18 Hypersil GOLD column (50 mm × 2.1 mm i.d., 3 µm particle size, Thermo Scientific, San Jose, CA, USA). A linear gradient was employed, using water (A) and methanol (B), both containing either 0.1 % formic acid. The gradient started with 20 % solvent B, increasing to 100 % over 1.5 minutes, holding for 1.5 minutes and returning to the initial 20 % B. The flow rate was maintained at 500 µL/min. The retention time of fentanyl was 1.41 min. The autosampler was set to 6°C, and the column was kept at room temperature. The HESI probe was operated in positive ion mode, with a spray voltage of 3.75 kV. The capillary temperature was set to 275°C, while the vaporizer temperature was 350°C. Nitrogen was used as the sheath gas (60 arbitrary units) and auxiliary gas (2 arbitrary units), and argon was used as the collision-induced dissociation (CID) gas at 1.5 mTorr.

Optimal collision energy (CE) for each transition was selected automatically. Quantification was performed by monitoring characteristic multiple reaction transition of m/z 337.2 ➔ 188.1 at 26 eV using internal standard method with a generic IS.

Data analysis included determining the internal fentanyl concentration and calculating percentage uptake (%) using the formula: (internal concentration/ external concentration) * 100. Results were visualized as a line graph comparing internal and external concentrations, generated using R, and the code, along with the raw data, is available on OSF.

### Light Sheet Microscopy

Light sheet microscopy was performed using a custom-built system based on the OpenSPIM platform as previously described by Winter et al. (2017, 2021). Briefly, Tg (*elavl3:GCaMP6)* larvae were pre-exposed to 50 LM fentanyl or 0.5% (v/v) DMSO in a well of a 24-well plate, and then transferred to a well containing drug plus the anti-nicotinic neuromuscular blocker, tubocurarine (4 mM), until muscle tone was lost (∼1 minute) (*n* = 7 larvae per treatment). Following drug exposure, larvae were transferred into a mixture of drug plus tubocurarine in 1.4% Low Melting Point (LMP) agarose and drawn into a clear borosilicate glass capillary (940 LM internal diameter) plugged with 1.5% LMP agarose. Once mounted, larvae were positioned vertically (head down) in the light sheet microscope, and optimal sectioning was undertaken repeatedly in the horizontal plane from the dorsal to ventral surface of the larvae, capturing 10 equidistant Z-planes spanning a total depth of 220 Lm. Each 6-minute imaging session generated a 200-frame Z-stack at a temporal resolution of 1.875 s per full brain volume. The larvae’s health was confirmed by checking for a normal heart rate and blood flow after each run to ensure viability.

The light sheet microscope system used a 488 nm Argon laser (Melles Griot, Netherlands) for excitation and a 20x/0.5 NA objective lens (Olympus, UK) for imaging, with an intermediary 1x magnification. Emitted light was filtered through a 525/50 nm emission filter and a 495 nm bandpass filter (Chroma, Germany). A 5.5 MP Zyla sCMOS camera (Andor, Northern Ireland) was used to capture images at 30 frames per second, 640 × 540-pixel resolution, and 4 × 4 binning with a 40 ms exposure. The camera and the rotational axis were controlled using the μManager software with the OpenSPIM plugin.

### Light Sheet Microscopy Image Processing and Fluorescence Analysis

Imaging data were processed using a custom-built Python pipeline, integrating functions form libraries including Scikit-image (van der Walt et al., 2014), SciPy (Virtanen et al., 2020) and Scikit-Learn (Pedregosa et al., 2011), available on <u>GitLab</u>, previously described by Winter et al. (2017, 2021). The processing began with 3D shift correction, followed by down-sampling by averaging over 3 x 3 x 1 voxel blocks to improve computational efficiency. Baseline correction was applied to each voxel by estimating the baseline using a sliding minimum filter, which was the subtracted from the data.

To align the datasets, a labelled brain template (Harvard Z-brain Atlas, Randlett et al. (2015)) was registered to each image stack in 3D using key-point registration. Fourteen anatomical landmarks were manually selected in each series, and the optimal affine transformation was computed using the Umeyama algorithm to align these points with their counterparts in the reference brain. This allowed for the registration of 45 predefined Regions of Interest (ROIs) encompassing major brain structures. Registration accuracy was visually assessed by overlaying the reference brain on the datasets, and any misaligned images were reprocessed by adjusting the key-points. The accuracy of the registration was quantified by calculating the median distance between the selected reference points and their aligned counterparts.

For each ROI, fluorescence intensity data were extracted and summarized as the time-averaged mean, median, and standard deviation of voxel intensities. The median was used as the primary measure of central tendency due to its robustness against small spatial registration errors. Temporal peak analyses were conducted on activity profiles, where the data were smoothed using a Gaussian filter (sigma = 1.5) to reduce high-frequency noise while preserving important signal features. Peaks were identified by applying a threshold of twice the standard deviation above baseline, and only those lasting for more than two consecutive imaging cycles (>1.875 seconds) were considered significant. Parameters such as peak height, width, interval between peaks, area under the curve (AUC), and the total number of peaks were recorded for each subject.

Statistical comparisons between fentanyl-exposed larvae and vehicle-treated controls were made using median fluorescence intensity data for each ROI. Normality was confirmed for each brain region using the Shapiro-Wilk method and following this either a Kruskal-Wallis test or one-way ANOVA was performed with either Dunn’s or Dunnett’s multiple comparison testing (respectively), and the percentage change in florescence intensity (%ΔF/F) between treatment groups was calculated using the formula:

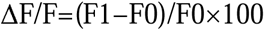

Where F1 represents fluorescence intensity of the fentanyl-treated group and F0 represents the corresponding mean value in the control group. Graphical representations were generated using GraphPad Prism 10 (GraphPad Prism, Version 10.2.3), Microsoft Excel and the custom Python script, with all code available on OSF and GitLab.

### Functional connectivity assessment

To measure functional connectivity, data were processed using a custom MatLab script. This script generated a connectivity matrix for each animal comprising Pearson’s correlation coefficients (r) between the time series’ of all possible pairs of ROIs. Next, post hoc statistical comparisons were undertaken between control and fentanyl-treated larvae using Mann– Whitney U tests with a Benjamini–Hochberg (BH) correction to compare the *r* values for each functional connection. A difference between these two values suggested an increase (positive) or decrease (negative) in the strength of that connection between untreated and fentanyl-treated brain states.

## Acknowledgements

We would like to thank the husbandry teams at the University of Portsmouth and University of Exeter for their support in rearing and care for the zebrafish.

## Author Contributions

Conceptualization, C.H., J.K. and M.O.P.; Methodology, C.H., J.K., M.T., M.J.W. and M.O.P.; Software, C.H. and M.J.W.; Validation, C.H. and M.T.; Formal Analysis, C.H., G.W. and M.T.; Resources, J.K., M.J.W. and M.O.P.; Data Curation, C.H., and M.T.; Writing – Original Draft, C.H.; Writing – Review & Editing, C.H., J.K., G.W., M.T., M.J.W. and M.O.P.; Visualization, C.H.; Supervision, J.K., M.J.W. and M.O.P.; Project Administration, J.K., M.J.W. and M.O.P.; Funding Acquisition, J.K., M.J.W. and M.O.P.

## Data Availability

All code and raw data files are available on OSF: https://osf.io/2jcts/?view_only=d4f08d1b813d410c8abfd1aa28768a40

## Ethical Statement

All experiments were approved by the University of Surrey and Exeter Animal Welfare and Ethical Review Board and under license from the UK Home Office.

## Supplementary Materials

### Bioavailability of fentanyl

To ensure that the observed locomotor responses were due to fentanyl uptake by the larvae following immersion exposure, we used LC-MS/MS to confirm the biological availability of fentanyl after immersion exposure to four fentanyl concentrations (2, 5, 10, and 20 μM) in 4 dpf larvae (**Supplementary Figure 1**). Previous research has shown that the larvae produce significantly different locomotor responses following addition of alarm chemicals to the well plate, thought to be because they perceive this as a threat (Wisenden et al., 2022). Therefore, here we also wanted to confirm that the observed behaviors are in all likelihood due to the systemic effects of the fentanyl, rather than resulting from a non-specific chemical effect, for example through olfactory detection. Although previous research has confirmed the uptake and biotransformation of fentanyl in 5 dpf larvae (Pesavento et al., 2022), no such information is available for 4 dpf larvae. Our bioanalysis revealed that all tested fentanyl concentrations were detected within the larvae at concentrations ranging from 130 to 170% of the external concentration. These data suggested sufficient uptake into the larvae following exposure and suggested a linear relationship between internal and external concentrations over the concentration range tested (**Supplementary Figure 1**). Although we were able to confirm compound uptake, due to the small tissue amounts available, it was not possible to measure tissue-specific levels of fentanyl and, as such, the actual concentration in the brain is not known.

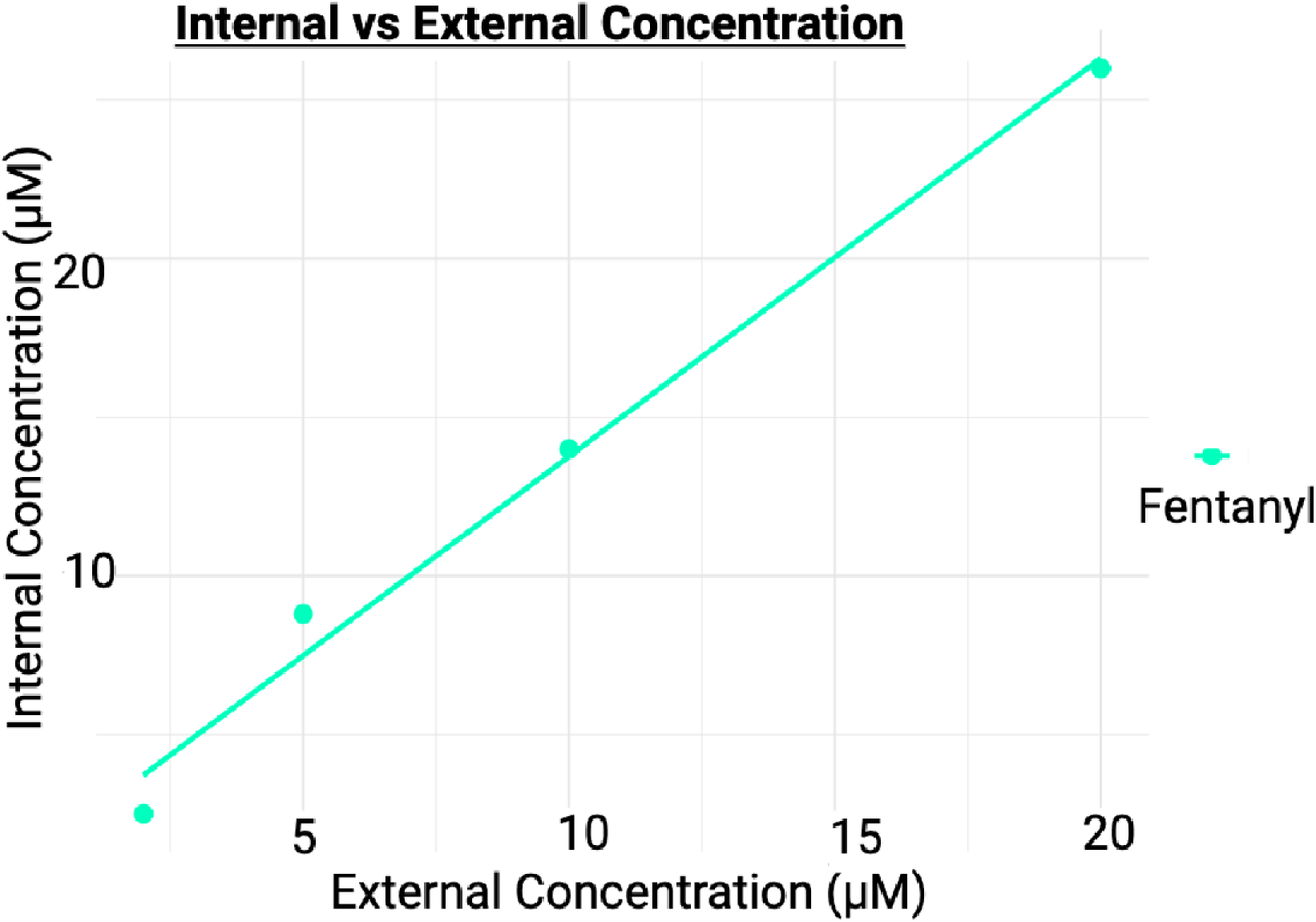

### The bioanalysis results for fentanyl exposure in 4 dpf zebrafish larvae

Bioanalysis was performed on larvae (*n* = 4) exposed to four different concentrations (2, 5, 10, 20 μM) of fentanyl to determine internal concentrations after exposure. The results show a linear relationship between internal concentration and external (exposed) concentration of fentanyl. Overall, these data are suggestive of a degree of bioaccumulation.

## References

Ables, J. L., Park, K., & Ibañez–Tallon, I. (2023). Understanding the habenula: A major node in circuits regulating emotion and motivation. Pharmacological Research, 190, 106734. 10.1016/j.phrs.2023.106734

Animals (Scientific Procedures) Act 1986, c. 14 (1986). https://www.legislation.gov.uk/ukpga/1986/14/contents

Arendt, F. (2021). The opioid-overdose crisis and fentanyl: The role of online information seeking via internet search engines. Health Communication, 36(10), 1148–1154. 10.1080/10410236.2020.1748820

Bae, Y.-K., Kani, S., Shimizu, T., Tanabe, K., Nojima, H., Kimura, Y., Higashijima, S.-I., & Hibi, M. (2009). Anatomy of zebrafish cerebellum and screen for mutations affecting its development. Developmental Biology, 330(2), 406–426. 10.1016/j.ydbio.2009.04.013

Chen, B.-Y., Jin, W.-Q., Chen, X.-J., Zhu, Y.-C., & Chi, Z.-Q. (2001). Analgesic activity and opioid receptor selectivity of stereoisomers of ohmefentanyl isothiocyanate. European Journal of Pharmacology, 424(3), 195–198. 10.1016/S0014-2999(01)01172-4

Choi, S. H., Koo, B.-N., Nam, S. H., Lee, S. J., Kim, K. J., Kil, H. K., Lee, K.-Y., & Jeon, D. H. (2008). Comparison of remifentanil and fentanyl for postoperative pain control after abdominal hysterectomy. Yonsei Medical Journal, 49(2), 204–210. 10.3349/ymj.2008.49.2.204

Colwill, R. M., & Creton, R. (2011). Locomotor behaviors in zebrafish (*Danio rerio*) larvae. Behavioural Processes, 86(2), 222–229. 10.1016/j.beproc.2010.12.003

Cooman, T., Bergeron, S. A., Coltogirone, R., Horstick, E., & Arroyo, L. (2021). Evaluation of fentanyl toxicity and metabolism using a zebrafish model. Journal of Applied Toxicology, 42(4), 706–714. 10.1002/JAT.4253

David, V., Durkin, T. P., & Cazala, P. (2002). Differential effects of the dopamine D2/D3 receptor antagonist sulpiride on self-administration of morphine into the ventral tegmental area or the nucleus accumbens. Psychopharmacology, 160(3), 307–317. 10.1007/s00213-001-0981-2

Duthie, D. J. (1998). Remifentanil and tramadol. British Journal of Anaesthesia, 81(1), 51–57. 10.1093/bja/81.1.51

Eastlack, S. C., Cornett, E. M., & Kaye, A. D. (2020). Kratom—pharmacology, clinical implications, and outlook: A comprehensive review. Pain and Therapy, 9(1), 55–69. 10.1007/s40122-020-00151-x

Edwards, C. J., & Kelley, D. B. (2001). Auditory and lateral line inputs to the midbrain of an aquatic anuran: Neuroanatomic studies in *Xenopus laevis*. Journal of Comparative Neurology, 438(2), 148–162. 10.1002/cne.1306

Faria, J., Barbosa, J., Moreira, R., Queirós, O., Carvalho, F., & Dinis-Oliveira, R. J. (2018). Comparative pharmacology and toxicology of tramadol and tapentadol. European Journal of Pain, 22(5), 827–844. 10.1002/EJP.1196

Fore, S., Palumbo, F., Pelgrims, R., & Yaksi, E. (2018). Information processing in the vertebrate habenula. Seminars in Cell & Developmental Biology, 78, 130–139. 10.1016/j.semcdb.2017.08.019

Giné, C. V., Espinosa, I. F., & Vilamala, M. V. (2014). New psychoactive substances as adulterants of controlled drugs. A worrying phenomenon? Drug Testing and Analysis, 6(7–8), 819–824. 10.1002/dta.1610

Gorwood, P. (2008). Neurobiological mechanisms of anhedonia. Dialogues in Clinical Neuroscience, 10(3), 291–299. 10.31887/DCNS.2008.10.3/pgorwood

Haehnel-Taguchi, M., Fernandes, A. M., Böhler, M., Schmitt, I., Tittel, L., & Driever, W. (2018). Projections of the diencephalospinal dopaminergic system to peripheral sense organs in larval zebrafish (*Danio rerio*). Frontiers in Neuroanatomy, 12, 20. 10.3389/fnana.2018.00020

Higashijima, S., Hotta, Y., & Okamoto, H. (2000). Visualization of cranial motor neurons in live transgenic zebrafish expressing green fluorescent protein under the control of the *Islet-1* promoter/enhancer. The Journal of Neuroscience, 20(1), 206–218. 10.1523/JNEUROSCI.20-01-00206.2000

Hill, R., Santhakumar, R., Dewey, W., Kelly, E., & Henderson, G. (2020). Fentanyl depression of respiration: Comparison with heroin and morphine. British Journal of Pharmacology, 177(2), 254–265. 10.1111/bph.14860

Hillman, C., Kearn, J., & Parker, M. O. (2024a). A unified approach to investigating 4 dpf zebrafish larval behaviour through a standardised light/dark assay. Progress in Neuro-Psychopharmacology and Biological Psychiatry, 134, 111084. 10.1016/j.pnpbp.2024.111084

Hillman, C., Kearn, J., & Parker, M. O. (2024b). Protocol for investigating light/dark locomotion in larval stage zebrafish using a standardized behavioral assay. STAR Protocols, 5(4), 103346. 10.1016/j.xpro.2024.103346

Hoyberghs, J., Bars, C., Ayuso, M., Van Ginneken, C., Foubert, K., & Van Cruchten, S. (2021). DMSO concentrations up to 1% are safe to be used in the zebrafish embryo developmental toxicity assay. Frontiers in Toxicology, 3, 804033. 10.3389/ftox.2021.804033

Kirla, K. T., Erhart, C., Groh, K. J., Stadnicka-Michalak, J., Eggen, Rik. I. L., Schirmer, K., & Kraemer, T. (2021). Zebrafish early life stages as alternative model to study ‘designer drugs’: Concordance with mammals in response to opioids. Toxicology and Applied Pharmacology, 419, 115483. 10.1016/j.taap.2021.115483

Knogler, L. D., Markov, D. A., Dragomir, E. I., Štih, V., & Portugues, R. (2017). Sensorimotor representations in cerebellar granule cells in larval zebrafish are dense, spatially organized, and non-temporally patterned. Current Biology, 27(9), 1288–1302. 10.1016/j.cub.2017.03.029

Komatsu, R., Turan, A. M., OrhanLSungur, M., McGuire, J., Radke, O. C., & Apfel, C. C. (2007). Remifentanil for general anaesthesia: a systematic review. Anaesthesia, 62(12), 1266–1280. 10.1111/j.1365-2044.2007.05221.x

Koyyalagunta, D. (2006). Opioid Analgesics. In Pain Management (Vol. 2, pp. 939–964). Elsevier. 10.1016/B978-0-7216-0334-6.50117-5

Lawson, R. P., Seymour, B., Loh, E., Lutti, A., Dolan, R. J., Dayan, P., Weiskopf, N., & Roiser, J. P. (2014). The habenula encodes negative motivational value associated with primary punishment in humans. Proceedings of the National Academy of Sciences, 111(32), 11858–11863. 10.1073/pnas.1323586111

Le Merrer, J., Becker, J. A. J., Befort, K., & Kieffer, B. L. (2009). Reward processing by the opioid system in the brain. Physiological Reviews, 89(4), 1379–1412. 10.1152/physrev.00005.2009

Le Merrer, J., Gavello-baudy, S., Galey, D., & Cazala, P. (2007). Morphine self-administration into the lateral septum depends on dopaminergic mechanisms: Evidence from pharmacology and Fos neuroimaging. Behavioural Brain Research, 180(2), 203–217. 10.1016/j.bbr.2007.03.014

Li, S., Guo, Y., Takahashi, M., Suzuki, H., Kosaki, K., & Ohshima, T. (2024). Forebrain commissure formation in zebrafish embryo requires the binding of KLC1 to CRMP2. Developmental Neurobiology, 84(3), 203–216. 10.1002/dneu.22948

Machluf, Y., Gutnick, A., & Levkowitz, G. (2011). Development of the zebrafish hypothalamus. Annals of the New York Academy of Sciences, 1220(1), 93–105. 10.1111/j.1749-6632.2010.05945.x

Metzger, M., Souza, R., Lima, L. B., Bueno, D., Gonçalves, L., Sego, C., Donato, J., & ShammahLLagnado, S. J. (2021). Habenular connections with the dopaminergic and serotonergic system and their role in stressLrelated psychiatric disorders. European Journal of Neuroscience, 53(1), 65–88. 10.1111/ejn.14647

Morgenstern, R., & Fink, H. (1985). Sulpiride blocks postsynaptic dopamine receptors in the nucleus accumbens. Journal of Neural Transmission, 61(3–4), 151–160. 10.1007/BF01251909

Nakazato, T., Horikawa, H. P. M., & Akiyama, A. (1998). The dopamine D2 receptor antagonist sulpiride causes long-lasting serotonin release. European Journal of Pharmacology, 363(1), 29–34. 10.1016/S0014-2999(98)00796-1

National Center for Biotechnology Information. (2024a). Naloxone. PubChem Compound Summary for CID 5284596. https://pubchem.ncbi.nlm.nih.gov/compound/Naloxone

National Center for Biotechnology Information. (2024b). PubChem Compound Summary for CID 261501, Tricaine Methanesulfonate. National Center for Biotechnology Information.

National Center for Biotechnology Information. (2024c). Remifentanil. PubChem Compound Summary for CID 60815. https://pubchem.ncbi.nlm.nih.gov/compound/Remifentanil

National Center for Biotechnology Information. (2024d). Sulpiride (Compound). National Center for Biotechnology Information.

National Center for Biotechnology Information. (2024e). PubChem Compound Summary for CID 3345, Fentanyl. National Center for Biotechnology Information. https://pubchem.ncbi.nlm.nih.gov/compound/Fentanyl

National Center for Biotechnology Information. (2024f). PubChem Compound Summary for CID 5462328, Heroin. National Center for Biotechnology Information. https://pubchem.ncbi.nlm.nih.gov/compound/Heroin

National Institute on Drug Abuse. (2021). Opioid Overdose Crisis. National Institutes of Health. https://www.drugabuse.gov/drug-topics/opioids/opioid-overdose-crisis

Nóbrega, L., & Dinis-Oliveira, R. J. (2018). The synthetic cathinone α-pyrrolidinovalerophenone (α-PVP): Pharmacokinetic and pharmacodynamic clinical and forensic aspects. Drug Metabolism Reviews, 50(2), 125–139. 10.1080/03602532.2018.1448867

Palieri, V., Paoli, E., Wu, Y. K., Haesemeyer, M., Grunwald Kadow, I. C., & Portugues, R. (2024). The preoptic area and dorsal habenula jointly support homeostatic navigation in larval zebrafish. Current Biology, 34(3), 489–504.e7. 10.1016/j.cub.2023.12.030

Pedregosa, F., Varoquaux, G., Gramfort, A., Michel, V., Thirion, B., Grisel, O., Blondel, M., Prettenhofer, P., Weiss, R., Dubourg, V., Vanderplas, J., Passos, A., Cournapeau, D., Brucher, M., Perrot, M., & Duchesnay, É. (2011). Scikit-learn: Machine learning in Python. Journal of Machine Learning Research, 12(85), 2825–2830.

Pesavento, S., Bilel, S., Murari, M., Gottardo, R., Arfè, R., Tirri, M., Panato, A., Tagliaro, F., & Marti, M. (2022). Zebrafish larvae: A new model to study behavioural effects and metabolism of fentanyl, in comparison to a traditional mice model. *Medicine*, Science and the Law, 62(3), 188–198. 10.1177/00258024221074568

Pose-Méndez, S., Schramm, P., Valishetti, K., & Köster, R. W. (2023). Development, circuitry, and function of the zebrafish cerebellum. Cellular and Molecular Life Sciences, 80(8), 227. 10.1007/s00018-023-04879-5

Proulx, C. D., Hikosaka, O., & Malinow, R. (2014). Reward processing by the lateral habenula in normal and depressive behaviors. Nature Neuroscience, 17(9), 1146–1152. 10.1038/nn.3779

Randlett, O., Wee, C. L., Naumann, E. A., Nnaemeka, O., Schoppik, D., Fitzgerald, J. E., Portugues, R., Lacoste, A. M. B., Riegler, C., Engert, F., & Schier, A. F. (2015). Whole-brain activity mapping onto a zebrafish brain atlas. Nature Methods, 12(11), 1039–1046. 10.1038/nmeth.3581

Rees, H., & Roberts, M. H. T. (1993). The anterior pretectal nucleus: a proposed role in sensory processing. Pain, 53(2), 121–135. 10.1016/0304-3959(93)90072-W

Rodda, L. N., Pilgrim, J. L., Di Rago, M., Crump, K., Gerostamoulos, D., & Drummer, O. H. (2017). A cluster of fentanyl-laced heroin deaths in 2015 in Melbourne, Australia. Journal of Analytical Toxicology, 41(4), 318–324. 10.1093/jat/bkx013

Sales Cadena, M. R., Cadena, P. G., Watson, M. R., Sarmah, S., Boehm li, S. L., & Marrs, J. A. (2021). Zebrafish (*Danio rerio*) larvae show behavioral and embryonic development defects when exposed to opioids at embryo stage. Neurotoxicology and Teratology, 85, 106964. 10.1016/J.NTT.2021.106964

Shafi, A., Berry, A. J., Sumnall, H., Wood, D. M., & Tracy, D. K. (2020). New psychoactive substances: A review and updates. Therapeutic Advances in Psychopharmacology, 10. 10.1177/2045125320967197

Stephenson-Jones, M., Floros, O., Robertson, B., & Grillner, S. (2012). Evolutionary conservation of the habenular nuclei and their circuitry controlling the dopamine and 5-hydroxytryptophan (5-HT) systems. Proceedings of the National Academy of Sciences, 109(3), E164–173. 10.1073/pnas.1119348109

Stewart, A. M., Braubach, O., Spitsbergen, J., Gerlai, R., & Kalueff, A. V. (2014). Zebrafish models for translational neuroscience research: From tank to bedside. Trends in Neurosciences, 37(5), 264–278. 10.1016/J.TINS.2014.02.011

Stewart, A. M., Gaikwad, S., Kyzar, E., & Kalueff, A. V. (2012). Understanding spatio-temporal strategies of adult zebrafish exploration in the open field test. Brain Research, 1451, 44–52. 10.1016/J.BRAINRES.2012.02.064

Stewart, A. M., & Kalueff, A. V. (2014). The behavioral effects of acute Δ9-tetrahydrocannabinol and heroin (diacetylmorphine) exposure in adult zebrafish. Brain Research, 1543, 109–119. 10.1016/J.BRAINRES.2013.11.002

Stewart, A., Wong, K., Cachat, J., Gaikwad, S., Kyzar, E., Wu, N., Hart, P., Piet, V., Utterback, E., Elegante, M., Tien, D., & Kalueff, A. V. (2011). Zebrafish models to study drug abuse-related phenotypes. Reviews in the Neurosciences, 22(1), 95–105. 10.1515/RNS.2011.011

Substance Abuse and Mental Health Services Administration. (2021). 2021 *National Survey on Drug Use and Health*. https://www.samhsa.gov/data/sites/default/files/reports/rpt39443/2021NSDUHFFRRev0 10323.pdf

Sutcliffe, K. J., Corey, R. A., Alhosan, N., Cavallo, D., Groom, S., Santiago, M., Bailey, C., Charlton, S. J., Sessions, R. B., Henderson, G., & Kelly, E. (2022). Interaction with the lipid membrane influences fentanyl pharmacology. Advances in Drug and Alcohol Research, 2, 10280. 10.3389/adar.2022.10280

Swanson, D. M., Hair, L. S., Strauch Rivers, S. R., Smyth, B. C., Brogan, S. C., Ventoso, A. D., Vaccaro, S. L., & Pearson, J. M. (2017). Fatalities involving carfentanil and furanyl fentanyl: Two case reports. Journal of Analytical Toxicology, 41(6), 498–502. 10.1093/jat/bkx037

Thompson, J. P., & Rowbotham, D. J. (1996). Remifentanil - an opioid for the 21st century. British Journal of Anaesthesia, 76(3), 341–343. 10.1093/bja/76.3.341

Tsuneoka, Y., & Funato, H. (2021). Cellular composition of the preoptic area regulating sleep, parental, and sexual behavior. Frontiers in Neuroscience, 15, 649159. 10.3389/fnins.2021.649159

Upadhyay, J., Maleki, N., Potter, J., Elman, I., Rudrauf, D., Knudsen, J., Wallin, D., Pendse, G., McDonald, L., Griffin, M., Anderson, J., Nutile, L., Renshaw, P., Weiss, R., Becerra, L., & Borsook, D. (2010). Alterations in brain structure and functional connectivity in prescription opioid-dependent patients. Brain, 133(Pt 7), 2098–2114. 10.1093/brain/awq138

van der Walt, S., Schönberger, J. L., Nunez-Iglesias, J., Boulogne, F., Warner, J. D., Yager, N., Gouillart, E., & Yu, T. (2014). scikit-image: image processing in Python. PeerJ, 2, e453. 10.7717/peerj.453

Vanwalleghem, G. C., Ahrens, M. B., & Scott, E. K. (2018). Integrative whole-brain neuroscience in larval zebrafish. Current Opinion in Neurobiology, 50, 136–145. 10.1016/j.conb.2018.02.004

Virtanen, P., Gommers, R., Oliphant, T. E., Haberland, M., Reddy, T., Cournapeau, D., Burovski, E., Peterson, P., Weckesser, W., Bright, J., van der Walt, S. J., Brett, M., Wilson, J., Millman, K. J., Mayorov, N., Nelson, A. R. J., Jones, E., Kern, R., Larson, E., … Vázquez-Baeza, Y. (2020). SciPy 1.0: Fundamental algorithms for scientific computing in Python. Nature Methods, 17(3), 261–272. 10.1038/s41592-019-0686-2

Vladimirov, N., Mu, Y., Kawashima, T., Bennett, D. V, Yang, C.-T., Looger, L. L., Keller, P. J., Freeman, J., & Ahrens, M. B. (2014). Light-sheet functional imaging in fictively behaving zebrafish. Nature Methods, 11(9), 883–884. 10.1038/nmeth.3040

Wang, B., Chen, J., Sheng, Z., Lian, W., Wu, Y., & Liu, M. (2022). Embryonic exposure to fentanyl induces behavioral changes and neurotoxicity in zebrafish larvae. PeerJ, 10, e14524. 10.7717/peerj.14524

Wang, H., Pélaprat, D., Roques, B. P., Vanhove, A., Chi, Z. Q., & Rostène, W. (1991). [3H]Ohmefentanyl preferentially binds to μ-opioid receptors but also labels σ-sites in rat brain sections. European Journal of Pharmacology, 193(3), 341–350. 10.1016/0014-2999(91)90149-K

Winter, M. J., Pinion, J., Tochwin, A., Takesono, A., Ball, J. S., Grabowski, P., Metz, J., Trznadel, M., Tse, K., Redfern, W. S., Hetheridge, M. J., Goodfellow, M., Randall, A. D., & Tyler, C. R. (2021). Functional brain imaging in larval zebrafish for characterising the effects of seizurogenic compounds acting via a range of pharmacological mechanisms. British Journal of Pharmacology, 178(13), 2671–2689. 10.1111/BPH.15458

Winter, M. J., Windell, D., Metz, J., Matthews, P., Pinion, J., Brown, J. T., Hetheridge, M. J., Ball, J. S., Owen, S. F., Redfern, W. S., Moger, J., Randall, A. D., & Tyler, C. R. (2017). 4-dimensional functional profiling in the convulsant-treated larval zebrafish brain. Scientific Reports, 7(1), 6581. 10.1038/s41598-017-06646-6

Wisenden, B. D., Paulson, D. C., & Orr, M. (2022). Zebrafish embryos hatch early in response to chemical and mechanical indicators of predation risk, resulting in underdeveloped swimming ability of hatchling larvae. Biology Open, 11(12). 10.1242/bio.059229

World Health Organization. (2021, August 4). Opioid overdose. World Health Organization. https://www.who.int/news-room/fact-sheets/detail/opioid-overdose

Zaig, S., Scarpellini, C., & Montandon, G. (2021). Respiratory depression and analgesia by opioid drugs in freely-behaving larval zebrafish. ELife, 10, e63407. 10.7554/ELIFE.63407

