## Supplementary for "Neural and Behavioral Dynamics of Acute Fentanyl Administration and Implications for Hazard Assessment of Novel Synthetic Opioids in Larval Zebrafish"

**Supplementary Materials**

**Bioavailability of fentanyl**

To ensure that the observed locomotor responses were due to fentanyl uptake by the larvae following immersion exposure, we used LC-MS/MS to confirm the biological availability of fentanyl after immersion exposure to four fentanyl concentrations (2, 5, 10, and 20 μM) in 4 dpf larvae (**Supplementary Figure 1**). Previous research has shown that the larvae produce significantly different locomotor responses following addition of alarm chemicals to the well plate, thought to be because they perceive this as a threat (Wisenden et al., 2022). Therefore, here we also wanted to confirm that the observed behaviors are in all likelihood due to the systemic effects of the fentanyl, rather than resulting from a non-specific chemical effect, for example through olfactory detection. Although previous research has confirmed the uptake and biotransformation of fentanyl in 5 dpf larvae (Pesavento et al., 2022), no such information is available for 4 dpf larvae. Our bioanalysis revealed that all tested fentanyl concentrations were detected within the larvae at concentrations ranging from 130 to 170% of the external concentration. These data suggested sufficient uptake into the larvae following exposure and suggested a linear relationship between internal and external concentrations over the concentration range tested (**Supplementary Figure 1**). Although we were able to confirm compound uptake, due to the small tissue amounts available, it was not possible to measure tissue-specific levels of fentanyl and, as such, the actual concentration in the brain is not known.

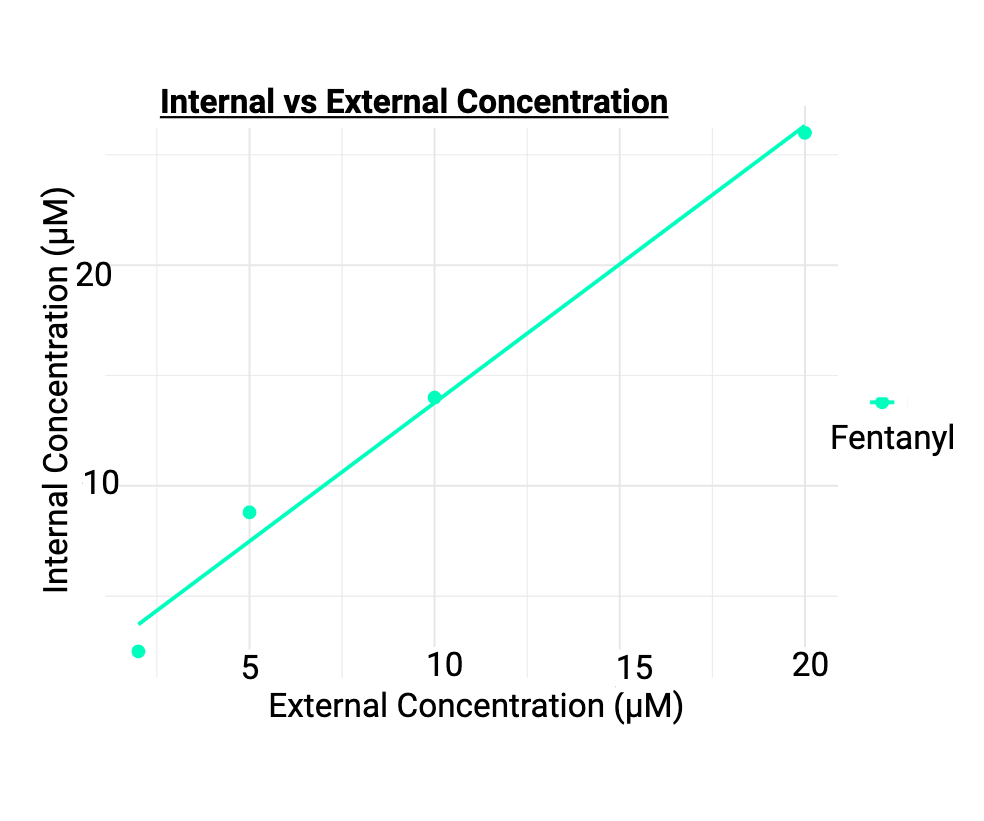

**The bioanalysis results for fentanyl exposure in 4 dpf zebrafish larvae.**

Bioanalysis was performed on larvae (*n* = 4) exposed to four different concentrations (2, 5, 10, 20 μM) of fentanyl to determine internal concentrations after exposure. The results show a linear relationship between internal concentration and external (exposed) concentration of fentanyl. Overall, these data are suggestive of a degree of bioaccumulation.
